# A simple method to determine the elimination half-life of drugs displaying noncumulative toxicity

**DOI:** 10.1101/761585

**Authors:** Deepesh Nagarajan, Preetham Venkatesh, Chandrani Thakur, Akshay Datey, Nagasuma Chandra, Dipshikha Chakravortty

**Affiliations:** Department of Biochemistry, Indian Institute of Science, Bangalore - 560012; Undergraduate program, Indian Institute of Science, Bangalore - 560012, India; Department of Microbiology and Cell Biology, Indian Institute of Science, Bangalore - 560012, India; Centre for Biosystems Science and Engineering, Indian Institute of Science, Bangalore - 560012, India

## Abstract

The pharmacokinetic characterization of a drug, especially the determination of its biological half-life, is an essential step during the early phases of drug development. An adequate half-life is amongst the many properties needed for selecting a drug candidate for clinical trials. Conversely, drug candidates possessing inadequate half-lives may be modified or eliminated from the drug discovery pipeline altogether. Several methods exist for determining the half-lives of drugs, namely HPLC, fluorescence assays, radioassays, radioimmunoassays, and elemental mass spectrometric assays. However, all these techniques are resource and labor-intensive, and cannot be used for the high-throughput half-life determination of hundreds of drug candidates. Here, we describe TOX_*HL*_: a simple technique to determine the half-lives of compounds displaying noncumulative toxicity. To calculate the half life, TOX_*HL*_ only relies on the survival outcomes of three experiments performed on an animal model: an acute toxicity experiment, a cumulative toxicity experiment, and a multi-dose experiment at different dosing intervals. As a proof of concept, we use TOX_*HL*_ to determine the peritoneal half-life of Ω76, an antimicrobial peptide. The half-life of Ω76 determined by TOX_*HL*_ is in good agreement with results from a standard mass spectrometric method, validating this approach.

## 1 Introduction

The biological half-life of a drug is the time required for half the drug to be eliminated from an organism, provided the rate of removal can be modeled as an exponential function^1^. Typically, the kidneys eliminate hydrophilic drugs from circulation, while the liver acts on hydrophobic compounds, adding polar groups for later renal excretion^2^. Drugs undergoing hepatic metabolism may also enter bile and be excreted through feces^3^. Proteolysis, the degradation of proteins into smaller peptides or amino acids, is also an important process for the elimination of therapeutic peptides^4^. Secondary drug removal also occurs through respiration, through the salivary glands^5^, mammary glands^6^, lacrimal glands^7^, skin^8^, and hair^9^.

A successful drug candidate should demonstrate adequate bioavailability^10^. The drug should remain at its site of action at a sufficient concentration, and for a sufficient time, for therapeutic effects to occur. During the early phases of drug development, measuring the half-life of a drug candidate in rodents is usually performed. Drug candidates possessing an adequate half-life for therapeutic effects to occur can enter later phases of development. Drug candidates possessing an insufficient half-life may need to be chemically altered and retested before entering later phases of development, or discarded altogether.

Numerous techniques exist for determining the concentrations and half-life of drugs. High performance liquid chromatography (HPLC) has been used to separate and characterize numerous small-molecule drugs based on molecular weights^11, 12^. Fluorescence assays can track a drug based on innate fluorescence or that of an attached fluorophore^13, 14^. Radioisotopes (typically ^14^C, but also ^32^P, ^34^S, etc.) incorporated into drug molecules allow them to be systemically tracked via scintillation counting^15^. Radioimmunoassays are competitive inhibition techniques where an unlabelled drug competes for antibody binding with its radioactive counterpart (typically labeled with ^3^H,^75^Se, ^125^I, etc.)^16–18^. Labeling a drug with a biologically rare element (such as Se) and tracking it using elemental mass spectrometry is also possible^19^. However, all these methods are labor-intensive, require sophisticated instruments, involve complicated chemical syntheses, or require handling radioisotopes. Such techniques are therefore impractical for the high-throughput screening of half-lives (100+ drugs at a time).

Here, we describe TOX_*HL*_: a simple method for the determination of the half-lives of drugs based on survival outcomes alone. TOX_*HL*_ involves 3 experiments: an acute toxicity experiment, a cumulative toxicity experiment, and a multi-dose experiment at different dosing intervals. Data obtained from these 3 experiments was sufficient to calculate the peritoneal half-life of an antimicrobial peptide (Ω76^19^) in BALB/c mice. The half-life of Ω76 calculated here was in good agreement with previous pharmacodynamic data obtained using mass spectrometry.

## 2 Results

### 2.1 Defining and determining (non)cumulative toxicity

Here, we define and contrast the terms *acute toxicity*, *maximum nonlethal dose*, *cumulative toxicity*, and *noncumulative toxicity*. We use two antimicrobial peptides: Ω76^19^ and Pexiganan^20, 21^, to demonstrate these concepts. Ω76 is a designed antimicrobial peptide effective against systemic infections of resistant *A. baumannii*. Here, Ω76 is used as an example drug that displays noncumulative toxicity and is compatible with TOX_*HL*_. Pexiganan is a broad-spectrum peptide that is effective at treating infected wounds. Here, Pexiganan is used as an example drug that displays cumulative toxicity and is incompatible with TOX_*HL*_. It should be noted that the toxicity results for Pexiganan described here should not be used to assess its suitability as a topical agent, a role where systemic toxicity is unimportant.

Acute toxicity comprises the lethal effects of a single dose of a drug, rather than the combined lethal effects of multiple doses of a drug. BALB/c mice (6-8 weeks, 20 g) were injected intraperitoneally with varying doses of Ω76 and Pexiganan, at 2-fold concentration increments. 5 mice per concentration cohort were used. All mice were observed for 5 days for mortality. Mice treated with both Ω76 and Pexiganan display mortality that increases in a dose-dependent manner. These results are depicted in Figure 1A. The LD_50_ values for Ω76 and Pexiganan were determined to be 170 mg/kg and 60 mg/kg respectively^19^, using linear (non-logarithmic) interpolation. However, for TOX_*HL*_, determining the *maximum nonlethal dose* (MND) rather than the LD_50_ is more important. We define the MND as the highest concentration of a drug, administered as a single intraperitoneal dose, that does not cause any mortality in mice observed for 5 days. Here, the MND values for Ω76 and Pexiganan are 32 mg/kg and 64 mg/kg respectively (Figure 1A, shown in arrows). These results are consistent with our previously reported values, which were performed on smaller cohorts^19^.

**Figure 1.**
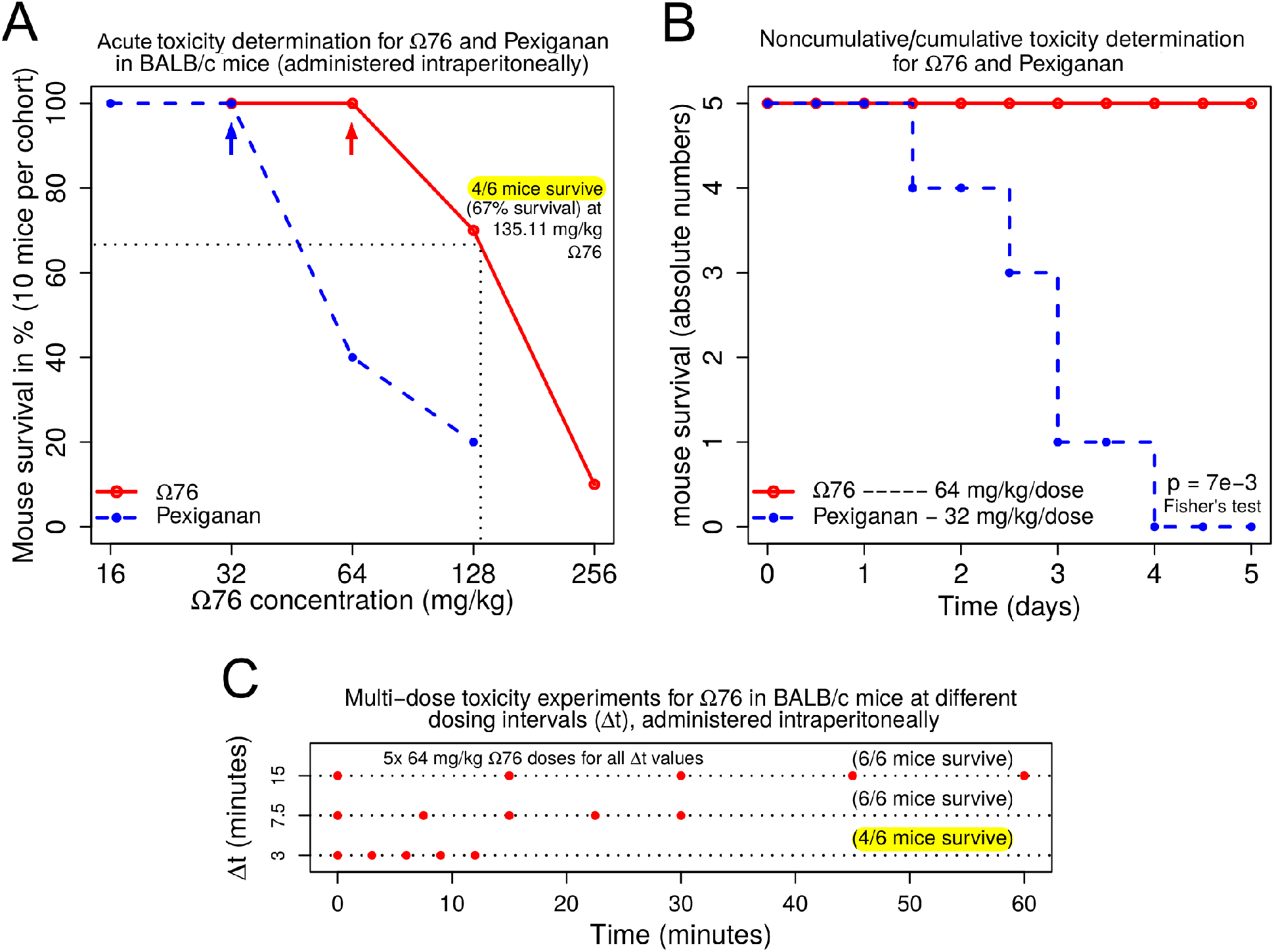
All experiments required to determine the elimination half-life using TOX_*HL*_. BALB/c mice (6-8 weeks, 20 g) were used for all experiments. **(A)** Acute toxicity experiments to determine survival vs. concentration of drug injected intraperitoneally. Determining the maximum nonlethal dose (MND), which is the highest concentration of a drug that causes no mortality, is essential for later noncumulative toxicity experiments. Here, the MND for Pexiganan is 32 mg/kg (blue arrow), and the MND for Ω76 is 64 mg/kg (red arrow). **(B)** Multi-dose (non)cumulative toxicity determination experiments. Here, mice were intraperitoneally injected with 11 doses of drug (at the MND concentration), at 12 hour intervals over the course of 5 days. All mice treated with Pexiganan died, indicating cumulative toxicity. All mice treated with Ω76 survived, indicating noncumulative toxicity. The difference between the survival outcomes of the Ω76 and Pexiganan cohorts are statistically significant (p=7e-3, Fisher’s test). **(C)** After Ω76 displayed noncumulative toxicity, we determined the dosing interval (Δt) between MNDs at which mortality is first observed. 5 doses at the MND were injected intraperitoneally at Δt of 15, 7.5 and 3 minutes. 67% survival (33% mortality) was observed a Δt = 3 minutes (Δt_*M*_, highlighted in yellow). The expected concentration of Ω76 responsible for 67% survival was calculated from panel **A** using linear interpolation (non-logarithmic), and found to be 135.11 mg/kg. Note that a separate set of similar experiments was described in our previous work^19^, but these experiments were only used to assess the toxicity of Ω76.

Cumulative toxicity comprises the lethal effects of multiple doses of drug when injected intraperitoneally at MND concentrations. 10 BALB/c mice (6-8 weeks, 20 g) were treated with 11 doses of Pexiganan, administered at the MND (32 mg/kg) at 12 hour intervals, over the course of 5 days. Despite Pexiganan never exceeding the MND concentration, 100% mortality was observed. This indicates that a single dose of Pexiganan causes a small amount of damage in mice, which is insufficient to cause mortality. However, this damage is not reversed within the dosing interval timespan (12 hours), and additional doses only serve to incrementally increase the total damage caused to mice, until a threshold is reached and death occurs. Autopsies indicate that the mechanism of cumulative toxicity for Pexiganan may involve cumulative damage to peritoneal organs culminating in intestinal inflammation (Figure 2B), which is absent in mice treated with Ω76 (Figure 2A).

**Figure 2.**
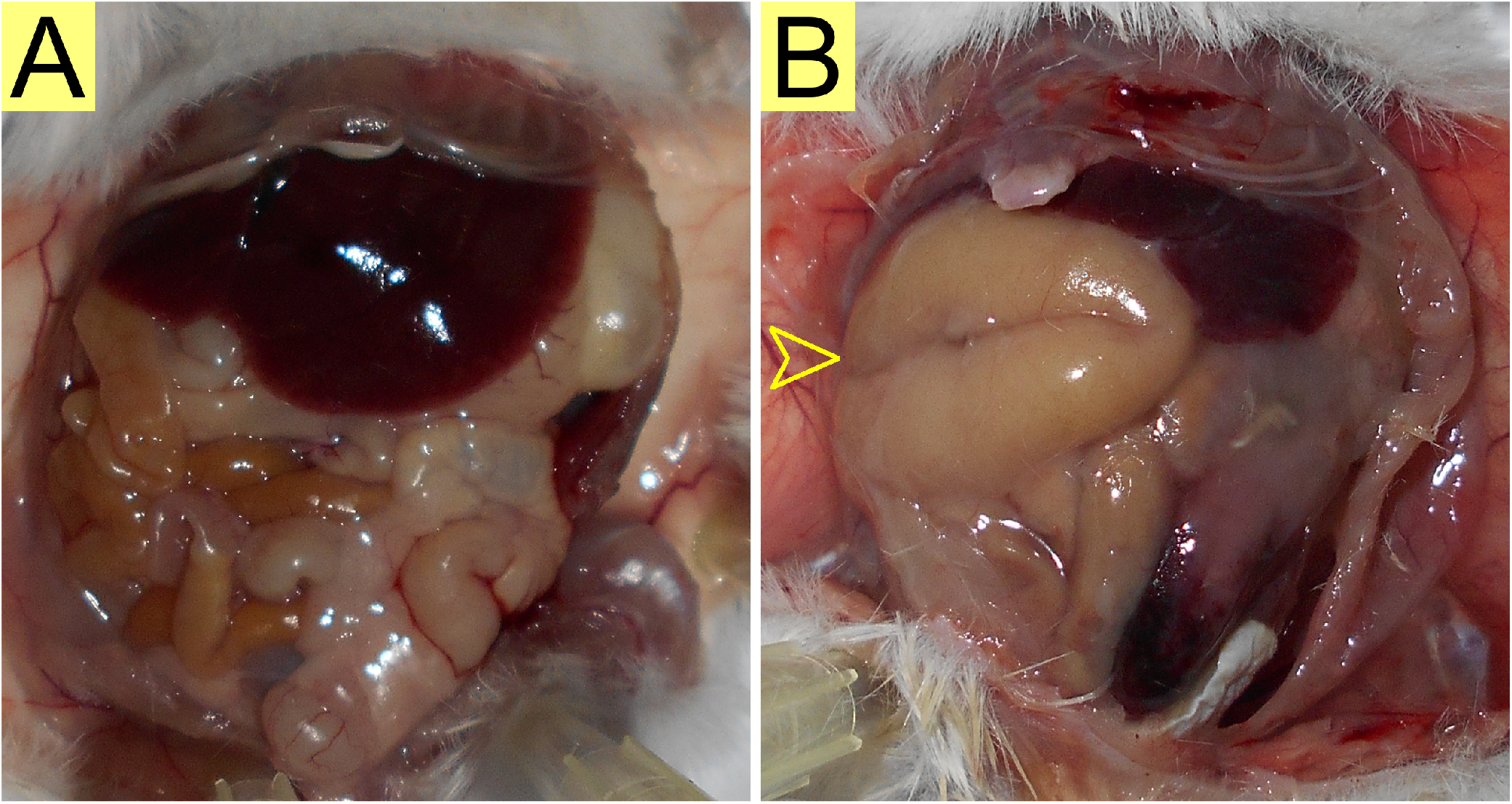
Peritonea of BALB/c mice (6-8 weeks, 20 g) treated with Ω76 and Pexiganan for (non)cumulative toxicity experiments. **(A)** Peritoneum of a mouse treated with 11 doses of 64 mg/kg Ω76 administered at 12 hour intervals over 5 days, and euthanized immediately after the last dose. All peritoneal organs appear intact. **(B)** Peritoneum of a mouse treated with 32 mg/kg Pexiganan administered at 12 hour intervals until mortality occurred. Here, intestinal inflammation is clearly visible (yellow arrow) and may be the cause of cumulative toxicity.

Noncumulative toxicity can be defined in contrast to cumulative toxicity. For a drug to display noncumulative toxicity, multiple doses injected intraperitoneally at MND concentrations should cause no mortality. 10 BALB/c mice (6-8 weeks, 20 g) were treated with 11 doses of Ω76, administered at the MND (64 mg/kg) at 12 hour intervals, over the course of 5 days. Ω76 never exceeded the MND concentration, and consequently neither intestinal inflammation nor mortality was observed (2A, Figure 1B). Our previous work^19^ further confirmed that Ω76 does not cause nephrotoxicity or hepatotoxicity in similarly treated mice. TOX_*HL*_ can only be used for drugs displaying noncumulative toxicity, which makes performing the acute and (non)cumulative toxicity experiments depicted in Figure 1A,B essential.

### 2.2 Experiments and input terms required for TOX_*HL*_

Once noncumulative toxicity for a given drug is established (Figure 1B), TOX_*HL*_ relies on the results of 2 experiments: acute toxicity experiments (Figure 1A) and multi-dose toxicity experiments (Figure 1C).

Multi-dose toxicity experiments are required to determine Δt_*M*_: the dosing interval that causes mortality, and 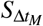: the percentage survival of mice at Δ*t*_*M*_. We determined the Δt_*M*_ by intraperitoneally injecting BALB/c mice (6-8 weeks, 20 g) with 5 doses of Ω76 at the MND (64 mg/kg) at different dosing intervals (Δt = 15, 7.5, and 3 minutes, Figure 1C). All mice were observed for 5 days. All mice survived Ω76 treatment for 5 days at Δt = 15 and 7.5 minutes. At Δ*t* = 3 minutes, only 4/6 mice (67%) survived for 5 days. Therefore, Δt_*M*_ = 3 minutes and 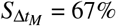. Note that unlike the *LD*_50_ or MND, Δt_*M*_ is not a constant for a given drug. Mortality would also be expected at all Δt ≤ 3 minutes. However, the ratio of 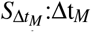 is expected to remain proportional, provided 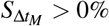. We recommend that a user must pick Δt values depending on the expected half life of the molecule being tested. If even the expected half life is unknown, a user can narrow down on the range using 2-fold Δt increases/decreases from an arbitrary starting Δt.

### 2.3 Using TOX_*HL*_ to calculate the elimination half-life

TOX_*HL*_ provides an estimate of *in vivo* half-life (*T*_1/2_) using only the values for MND, Δt_*M*_, 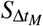, and the acute toxicity plot depicted in Figure 1A. TOX_*HL*_ begins by constructing a theoretical concentration vs. time model (Figure 3A) for the experiment used to determine Δt_*M*_ (Figure 1C). 5 Ω76 doses at the MND (64 mg/kg) were administered at Δt_*M*_ = 3 minutes. Ω76 absorption into the peritoneum is modeled as instantaneous, as peritoneal injections can be administered in 1-2 seconds. The peritoneal elimination of Ω76 is modeled as a first order (exponential) process. Over the course of 12 minutes (4 × Δt_*M*_), the Ω76 peritoneal concentration varies from 0 to N_8_, with intermediate concentrations ranging from *N*_1_ → *N*_7_. *N*_*k*_ represents a constant concentration increase corresponding to the MND of 64 mg/kg. For contrast, a concentration vs. time model for a larger dosing interval (Δ*t* = 15 minutes) is also provided (Figure 3B).

**Figure 3.**
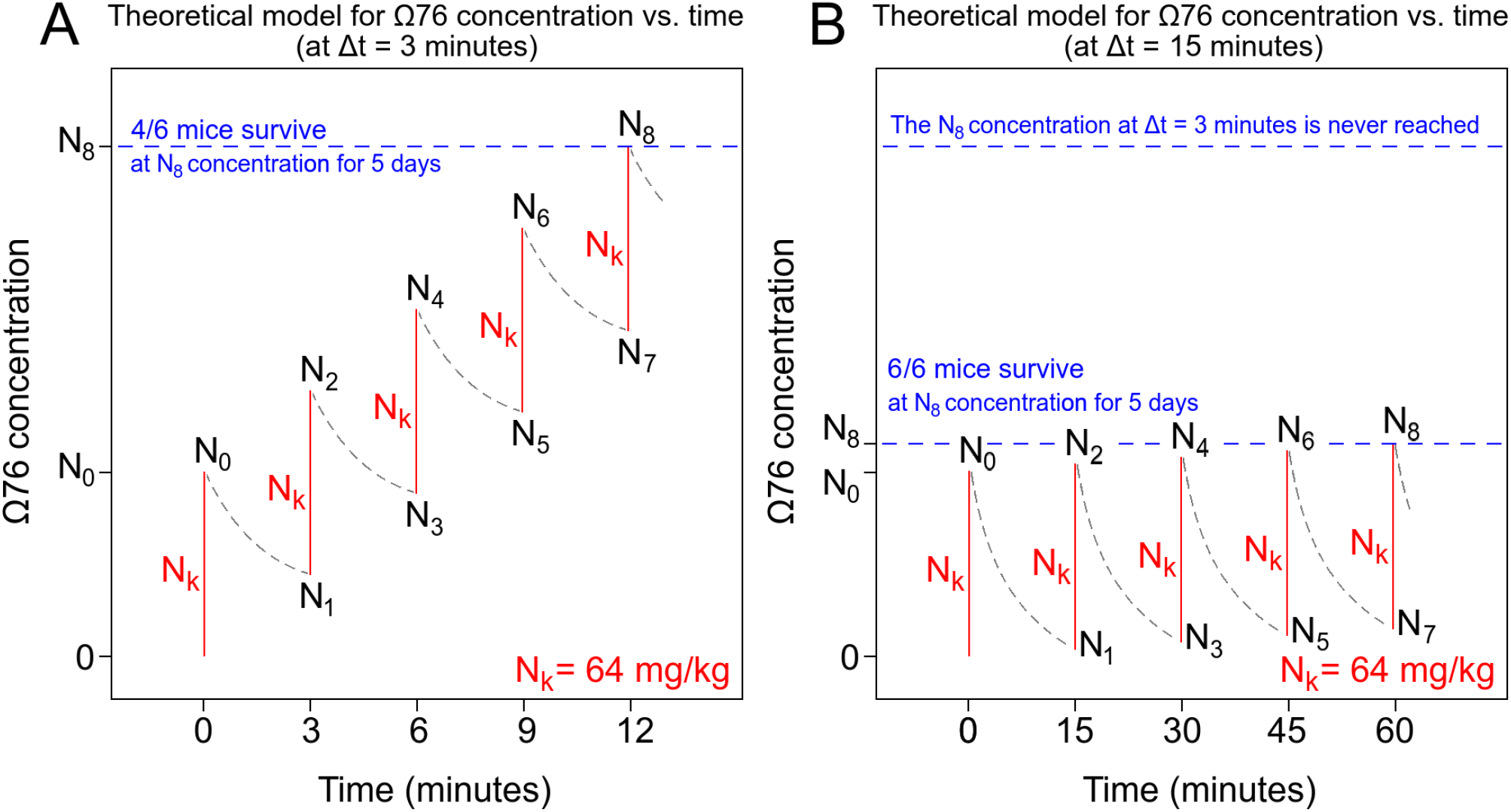
Theoretical pharmacokinetics of a multi-dose experiment involving administration of 5 doses of Ω76 at the MND (64 mg/kg, here denoted as *N*_*k*_). **(A)** At Δt = 3 minutes (previously described in Figure 1C). Ω76 absorption into the peritoneum is modeled as instantaneous (red lines), as intraperitoneal injections can be administered in 1-2 seconds. Ω76 elimination from the peritoneum is modeled using first order kinetics (exponential elimination, gray curves). From Figure 1A,C, we determined that the highest Ω76 concentration reached in the peritoneum (*N*_8_) was 135.11 mg/kg. Given Δt, *N*_*k*_, and *N*_8_, the values of *N*_0_ → *N*_8_, can easily be determined. These values would allow us to calculate the *in vivo* half-life of Ω76. **(B)** For contrast, the expected pharmacokinetics of Ω76 at Δt = 15 minutes is provided. Note that the concentration of Ω76 will not reach the previous *N*_8_ concentration (at Δt = 3 minutes) after 5 doses, and no mortality will occur.

Figure 3 can also be described mathematically, using Equation-set 1:

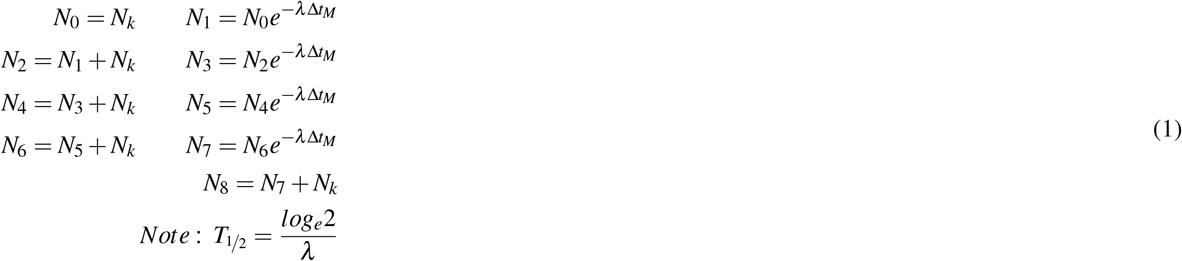

Equation-set 1 can be simplified as described in Equation-set 2.

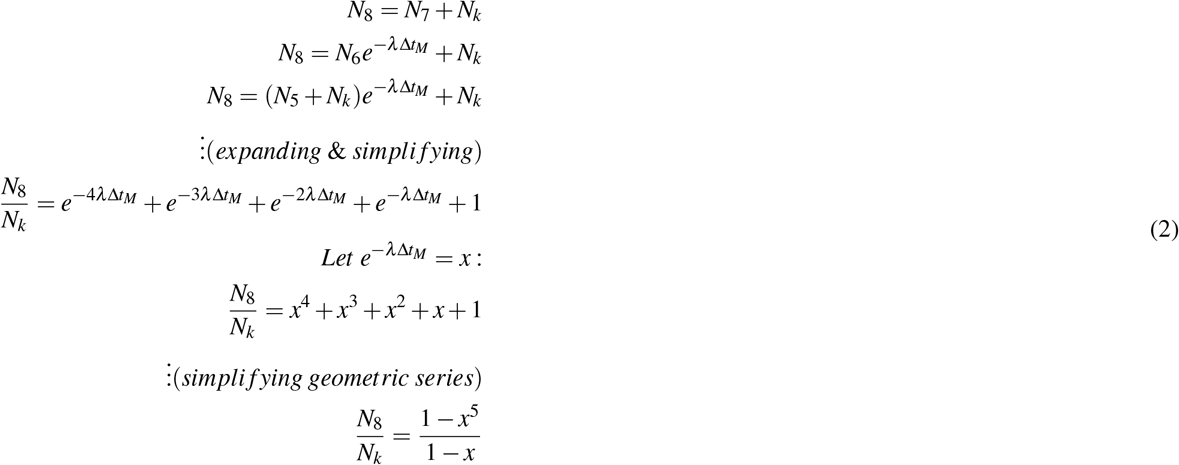

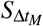 (67%) mice survive at concentration N_8_ (Figure 1C, highlighted). Using linear (non-logarithmic) interpolation, 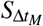 survival occurs at a single-dose Ω76 concentration of 135.11 mg/kg (Figure 1A, highlighted). Once the value of N_8_/N_*k*_ is known, Equation-set 2 can be solved for *x*. *T*_1/2_ can be calculated from *x* using Equation-set 3.

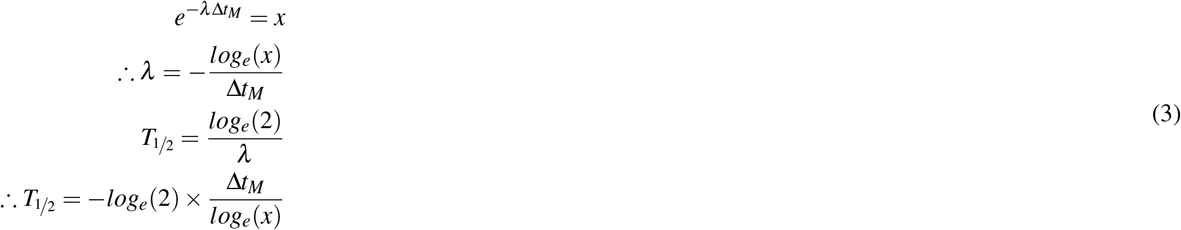

For Ω76, N_8_/N_*k*_ = 135.11/64 = 2.11, *x* = 0.55, and *T*_1/2_ = 3.48 minutes.

Note that the description of TOX_*HL*_ provided above describes equations for a 5-dose cumulative toxicity experiment (Figure 1C). The solution provided in Equation-set 2 can easily be modified for an *n*-dose cumulative toxicity experiment (Equation 4, where n ≥ 2).

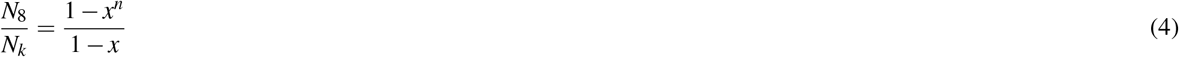

All experiments and calculations required to calculate *T*_1/2_ using for TOX_*HL*_ are provided in Figure 4. A python script that calculates *T*_1/2_ using TOX_*HL*_ is provided on GitHub (https://github.com/preetham-v/TOX_HL) and is made available as a webserver (http://proline.biochem.iisc.ernet.in/toxhl/).

**Figure 4.**
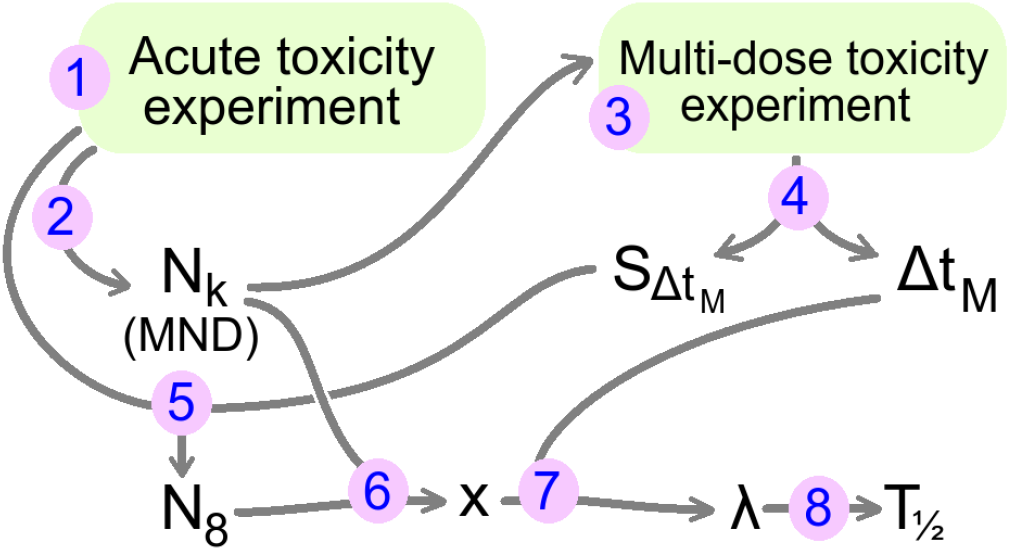
All experiments and calculations used by TOX_*HL*_ to calculate *T*_1/2_ are provided in order. (1): Perform the acute toxicity experiment (Figure 1A). (2): obtain *N*_*k*_ value from acute toxicity experiment. (3): Use *N*_*k*_ to guide the dosing concentration for the multi-dose toxicity experiment (Figure 1B). (4): Obtain values for Δ*t*_*M*_ and 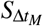 from the multi-dose toxicity experiment. (5): Calculate *N*_8_ concentration by looking at the mortality corresponding to 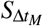 in the acute toxicity graph. (6): Calculate *x* using the values of *N*_*k*_ and *N*_8_, using Equation-set 2. (7,8): Use *x* to calculate *λ* and *T*_1/2_ using Equation-set 3.

### 2.4 Validating TOX_*HL*_ using data from a standard pharmacokinetic assay

Our previous work involved the pharmacokinetic characterization of Ω76 in mice^19^. 70 mg/kg N selenomethionine-labeled Ω76 (Nselmet-Ω76, the molar equivalent of 64 mg/kg unlabeled Ω76) was injected intraperitoneally into BALB/c mice (6-8 weeks, 20 g), and withdrawn at different time intervals using cardiac punctures. We obseved that Nselmet-Ω76 reached a serum concentration maxima (*C*_*max*_) at 4.59 minutes post-injection. This indicates that the peritoneal elimination half-life of Ω76 is within the range of 0 → 4.59 minutes, and is in good agreement with the TOX_*HL*_ value of 3.48 minutes.

## 3 Discussion

In this work, we described TOX_*HL*_, using which the half-life of a drug can be determined based on the survival outcomes of 3 simple experiments performed on mice. TOX_*HL*_ is cost-effective, easy to perform, and requires no instruments or special reagents. We described and validated TOX_*HL*_ using an example antimicrobial peptide (Ω76). Using TOX_*HL*_, the peritoneal half-life of Ω76 in mice was found to be 3.48 minutes. This value was in good agreement with the peritoneal half-life range of 0 → 4.59 minutes, calculated using a conventional ICP-MS technique that tracked selenium. A python script (https://github.com/preetham-v/TOX_HL) and webserver (http://proline.biochem.iisc.ernet.in/toxhl/) implementing TOX_*HL*_ have beem made available to the community.

TOX_*HL*_ can be modified in many ways to meet a user’s requirements. We have used 5 intraperitoneal doses (*n*) to determine _*M*_ and *S*_*M*_ (Figure 1C). However, any number of doses (2 ≤ *n* < ∞) may be used. *S*_*M*_ will increase with *n*, which is advantageous as it eliminates time-errors when dosing at a small. However, the amount of drug used will increase with *n*, necessitating the synthesis/purchase of greater quantities. The total volume of drug injected intraperitoneally will also increase. Extreme values of *n* (for example, 20 doses of 200 *μ*L each), would cause abdominal distention in mice, altering the elimination kinetics. Larger *n* should therefore be accompanied by smaller injection volumes. TOX_*HL*_ models first order elimination kinetics (3). For exceptional drugs following non-exponential kinetics, a user can replace the exponential equations in Equation-set 1 with decay equations of their choosing. TOX_*HL*_ can also be used to calculate the bloodstream half-life of drugs, simply by performing intravenous rather than intraperitoneal injections in mice.

It should be noted that TOX_*HL*_ was only validated using a single molecule (Ω76). Additional experiments using more molecules would help further validate our approach. TOX_*HL*_ can only be used for drugs possessing noncumulative toxicity, which has to be tested prior to half-life calculation (Figure 1B). TOX_*HL*_ requires a larger number of mice in comparison to conventional methods. Consequently, a larger amount of drug is required for these tests.

Nevertheless, we expect TOX_*HL*_ to be especially useful for the half-life determination of a library of compounds, created from a parent compound displaying noncumulative toxicity. For example, a 100-1000 member peptide library, created using saturation mutagenesis of a parent antimicrobial peptide, could quickly and easily be assayed using TOX_*HL*_.

## 4 Methods

For all experiments described in this work, BALB/c mice (6-8 weeks, 20 g) were housed at the Central Animal Facility (CAF, IISc) and fed *ad libitum* with pellet feed and water. All animal experiments were approved by the Institutional Animal Ethics Committee, IISc (Project No. CAF/Ethics/550/2017). A detailed description of all methods used is provided in the results section. However, it should be noted that 20 mg/mL stocks for both Ω76 and Pexiganan were prepared in physiological saline (0.8% NaCl) and stored at −80 °C. Both peptides display poorer solubility in phosphate buffered saline (PBS). These stocks were diluted in saline to obtain the concentration needed for the specific experiment. All intraperitoneal injections were 200 *μ*L in volume, as larger volumes could alter peritoneal elimination kinetics.

## Author contributions statement

D.N. conceived and designed TOX_*HL*_, and performed all experiments. P.V. created a python script and webserver implementing TOX_*HL*_. C.T. and A.D. co-performed mouse experiments. D.C. and N.C. coordinated the study, planned experiments, and provided resources. All authors reviewed the final draft of the manuscript.

